# miR2105 regulates ABA biosynthesis via OsbZIP86-*OsNCED3* module to contribute to drought tolerance in rice

**DOI:** 10.1101/2021.09.21.461241

**Authors:** Weiwei Gao, Mingkang Li, Songguang Yang, Chunzhi Gao, Yan Su, Xuan Zeng, Zhengli Jiao, Weijuan Xu, Mingyong Zhang, Kuaifei Xia

## Abstract

Induced abscisic acid (ABA) biosynthesis plays an important role in plant tolerance to abiotic stresses, including drought, cold and salinity. However, regulation pathway of the ABA biosynthesis in response to stresses is unclear. Here, we identified a rice miRNA, osa-miR2105 (miR2105), which plays a crucial role in ABA biosynthesis under drought stress. Analysis of expression, transgenic rice and cleavage site showed that *OsbZIP86* is a target gene of miR2105. Subcellular localization and luciferase activity assays showed that OsbZIP86 is a nuclear transcription factor. *In vivo* and *in vitro* analyses showed that OsbZIP86 directly binds to the promoter of *OsNCED3*, and interacts with OsSAPK10, resulting in enhanced-expression of *OsNCED3*. Transgenic rice plants with knock-down of miR2105 or overexpression of *OsbZIP86* showed higher ABA content, more tolerance to drought, a lower rate of water loss, more stomatal closure than wild type rice ZH11 under drought stress. These rice plants showed no penalty with respect to agronomic traits under normal conditions. By contrast, transgenic rice plants with miR2105 overexpression, *OsbZIP86* downregulation, or *OsbZIP86* knockout displayed less tolerance to drought stress and other phenotypes. Collectively, our results show that a regulatory network of ‘miR2105-OsSAPK10/OsbZIP86-*OsNCED3*’ control ABA biosynthesis in response to drought stress.

**One-sentence summary:** ‘miR2105-OsbZIP86-*OsNCED3*’ module plays crucial role in mediating ABA biosynthesis to contribute to drought tolerance with no penalty with respect to agronomic traits under normal conditions.

## Introduction

Rice (*Oryza sativa* L.) is one of the most important grain crops in the word, and both biotic and abiotic stresses are major threats to its yield. Therefore, breeding programs are directed towards increasing yield and improving tolerance to biotic and abiotic stresses (Kumar *et al*., 2018). To cope with most stresses, plants synthesize abscisic acid (ABA), which activates ABA-mediated signaling pathways, including stomatal closure under drought stress, metabolic adjustment, growth regulation and regulation of defense-related genes (Finkelstein *et al*., 2002; Joo *et al*., 2020). Therefore, ABA is one of the most important phytohormones regulating plant growth, development, and stress response (Nambara & Marion-Poll, 2005; Chen *et al*., 2020). The core components of ABA biosynthesis, catabolism, transport, and signaling have been identified (Chen *et al*., 2020). ABA is synthesized in the plastids and the cytosol from zeaxanthin in a five-step biosynthetic process in *Arabidopsis*. The first three steps of conversion from the precursor β-carotene to violaxanthin and neoxanthin, which are catalyzed by ABA DEFICIENT 1 (ABA1), ABA4, and 9-*cis*-epoxycarotenoid dioxygenases (NCEDs), take place in the plastids; violaxanthin and neoxanthin are then transported into the cytosol for the next two steps, in which ABA is produced by the catalysis of ABA2 and ABA3 in Arabidopsis (Nambara & Marion-Poll, 2005; Chen *et al*., 2020).

NCEDs cleave violaxanthin and neoxanthin to produce xanthoxin in the ABA biosynthetic pathway; this is the rate-limiting step of ABA *de novo* biosynthesis (Nambara & Marion-Poll, 2005; Dong *et al*., 2015). *NCED*s belong to a multigene family in plants, and their expression is tightly regulated in response to developmental or stress conditions. Rice has five *NCED*s. *OsNCED1* is most highly expressed in leaves as a housekeeping gene under normal conditions and is feedback-regulated by ABA, suppressed by water stress, and induced in cold-stressed anthers (Ye *et al*., 2011). *OsNCED2* has roles in the seed germination (Zhu *et al*., 2009) and grain ABA production (Nonhebel & Griffin, 2020). *OsNCED3, OsNCED4* and *OsNCED5* mediate ABA biosynthesis, and confer to different stress tolerance (Huang *et al*., 2018; Hwang *et al*., 2018; Huang *et al*., 2019). *OsNCED3* is constitutively expressed in various tissues under normal conditions, and responses to multi-abiotic stresses in plant growth (Huang *et al*., 2018). The expression of *OsNCED3* is rapidly induced by drought stress and quickly decreased after rehydration; thus, it is a major gene promoting ABA biosynthesis during drought stress in rice (Ye *et al*., 2011; Mao *et al*., 2017; Liu *et al*., 2018).

The 20–22 nucleotide microRNAs (miRNAs) have critical roles in both plant development and biotic and abiotic stress responses, including ABA response (Nadarajah & Kumar, 2019). miRNAs can regulate gene expression at post-transcriptional levels through specific base-pairing to target mRNAs (Bartel, 2004). In rice, a serial of miRNAs, such as miR156, miR159, miR168, miR169, miR319, and miR395 are involved in various stress response (Zhou *et al*., 2010). miR159 is induced by drought stress and provides tolerance against abiotic stresses such as drought in rice (Mohsenifard *et al*., 2017). miR319 is also induced by salinity and drought stress and makes a positive contribution to the plant abiotic stress response (Koyama *et al*., 2017). In Arabidopsis, miR165 and miR166 regulate expression of BG1, a glucosidase; BG1 hydrolyzes Glc-conjugated ABA, which in turn further modulates ABA homeostasis (Yan *et al*., 2016). miR2105 has been isolated from developing rice seeds (Xue *et al*., 2009), but its role has remained elusive.

The bZIP transcription factors (TFs) also have crucial regulatory roles in activating ABA-dependent stress-responsive gene expression (Joo *et al*., 2021). It is predicated that rice encodes 89 bZIP TFs, several of which have been found to be involved in rice stress responses (Joo *et al*., 2021). For example, OsbZIP23 positively regulates the transcription of *OsNCED4* to mediate ABA biosynthesis (Xiang *et al*., 2008; Zong *et al*., 2016). OsbZIP71 directly binds to G-box sequences in the promoters of *OsNHX1* and *COR413-TM1* to contribute to drought and salt tolerance (Liu *et al*., 2014). Rice OsbZIP86 (previously known as osZIP-1a) is a homolog of wheat G-box-binding factor EmBP-1; overexpression of *osZIP-1a* in rice protoplasts can enhance expression from the wheat *Em* gene promoter containing G-boxes only in the presence of ABA (Nantel & Quatrano, 1996). Moreover, phosphorylated OsbZIP72 directly binds to the G-box in the promoter of *AOC*, and activates *AOC* transcription (Wang *et al*., 2020).

Members of the phosphorylation-activated sucrose nonfermenting 1–related protein kinase 2 (SnRK2) have also been reported to have crucial roles in phosphorylating AREB/ABF TFs, and subsequently activating downstream genes to respond to ABA signals (Banerjee & Roychoudhury, 2017). The rice SnRK2 protein family contains 10 members, denoted stress/ABA-activated protein kinase 1 (SAPK1) to SAPK10 (Kobayashi *et al*., 2004). Among them, SAPK1, SAPK2, SAPK6, SAPK8, SAPK9, and SAPK10 have been reported to be functionally related to ABA signaling (Wang *et al*., 2020; Fu *et al*., 2021). SnRK2s usually function through their potential substrate proteins. A few TFs have been identified as SnRK2 substrates, including ABI5, OsbZIP23, OsbZIP46, OsbZIP62 and OsbZIP72 (Rehman *et al*., 2021).

Previous efforts have provided evidence that miRNAs and bZIP TFs contribute to ABA biosynthesis in regulating drought resistance (Nadarajah & Kumar, 2019). However, regulation pathway of ABA biosynthesis and the cross talk between miRNAs and target genes are not well understood in rice. Here, we report that *OsbZIP86*, the target gene of miR2105, regulates drought-induced ABA biosynthesis through directly increasing the expression of *OsNCED3* to modulate drought tolerance, without penalty with respect to main agronomic traits under normal conditions. OsSAPK10 can interact with OsbZIP86, and increase *OsNCED3* promoter activity. Taken together, our results demonstrate the crucial importance of the ‘miR2105-OsSAK10/OsbZIP86-*OsNCED3*’ module in mediating ABA biosynthesis under drought stress.

## Results

### *OsbZIP86* is a target gene of miR2105

Previously, we identified a drought-repressed miRNA osa-miR2105 (miR2105) from rice seedlings by miRNA sequencing; this miRNA was also isolated from developing rice seeds (Xue *et al*., 2009; Yi *et al*., 2013). In order to investigate the functions of miR2105 in stress responses in rice, we detected its expression under ABA, drought, and salt treatments by real-time qRT-PCR. The expression of miR2105 was repressed under ABA treatment in wild-type rice ZH11 seedlings (Figure **1A**). miR2105 was also downregulated after 0.5–4 h of water deficiency treatment and returned to relatively high expression levels after 1 h of re-watering (Figure **1B**). Expression of miR2105 was also downregulated by NaCl treatment (Figure **1C**). These results suggest that miR2105 is an ABA-, drought-, and salt-repressed miRNA.

**Figure 1.**
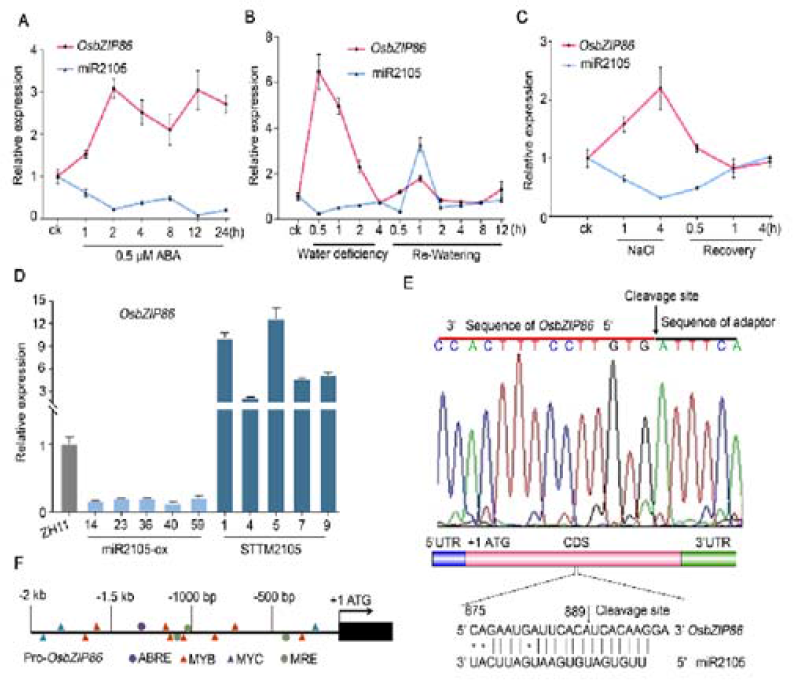
miR2105 regulates expression of *OsbZIP86* under ABA, drought, and salt treatments. (A–C) Expression changes of *OsbZIP86* and miR2105 under ABA, drought, and salt treatments. RNA was isolated from 2-week-old ZH11 rice seedlings grown in Yoshida solution supplied with 0.5 μM ABA at the indicated time (A). For drought and salt treatments, 2-week-old seedlings grown in Yoshida solution were exposed to air (B) or treated with 150 mM NaCl (C) for 4 h, and then the seedlings were transferred to Yoshida solution again for recovery. (D) Expression changes of *OsbZIP86* in transgenic rice overexpressing osa-miR2105 (miR2105-ox) or with downregulation of osa-miR2105 (STTM2105) under normal growth conditions. (E) The cleavage site targeted by miR2105 in the *OsbZIP86* mRNA. The arrow on the miRNA:mRNA alignment indicates the cleavage site from 10 sequencing clones identified in ZH11 seedlings by 5′-RLM-RACE. *U6* and *e-EF-1a* were used as miRNA and mRNA reference genes, respectively, and mean ± SD (n = 3) values are shown in (A–D). All qRT-PCR analyses for gene expression were performed in three biological replicates with similar results. (F) Main stress-related *cis*-acting elements in the 2-kb *OsbZIP86* promoter region. The *cis* elements are indicated.

It is well known that miRNAs exert their functions by inhibiting the expression of their target genes (Bartel, 2004). To identify the target gene of miR2105, we used two methods to identify its downstream targets. A total of 13 genes were predicted to be targets of miR2105 using psRNATarget (http://plantgrn.noble.org/psRNATarget/) (Supplemental Table **S1**). To further validate the targets, we generated miR2105-overexpressing (miR2105-ox) and miR2105-downregulation lines (STTM2105) (Supplemental Figure **S1A**,**B**). Among the predicted 13 target genes, *OsbZIP86* (LOC_Os12g13170) (Nijhawan *et al*., 2008) was downregulated in the T_2_–T_4_ generations of miR2105-ox and upregulated in the T_2_–T_4_ generations of STTM2105 (Figure **1D** and Supplemental Figure **S2A**). These results show that the expression change of *OsbZIP86* occurred in the opposite direction to that of miR2105 in miR2105-ox and STTM2105.

To further confirm *OsbZIP86* as a target gene of miR2105, a 5′ RNA ligase-mediated rapid amplification of cDNA ends (5′-RLM-RACE) assay was performed *in vivo* using ZH11 seedlings grown under normal condition. Sequencing of the 5′-RLM-RACE clones revealed that *OsbZIP86* mRNA was cleaved at the miR2105/*OsOsbZIP86* mRNA complementary site (Figure **1E**). We also searched the *OsbZIP86* expressed sequence tags (ESTs) in the NCBI database and found that an EST (CF324346) had its first base pair located within the complementary site. Collectively, these data demonstrated that miR2105 could direct cleavage of *OsbZIP86* mRNA to regulate transcript levels of *OsbZIP86*.

### Drought, salt and ABA induce expression of *OsbZIP86* and repress that of miR2105

*OsbZIP86* (*osZIP-1a*) belongs to the OsbZIP TF family (Nantel & Quatrano, 1996; Nijhawan *et al*., 2008), and its promoter contains putative stress response-related *cis*-elements (Figure **1F**). Through searching the Rice eFP database (http://bar.utoronto.ca/efprice/cgi-bin/efpWeb.cgi), we found that drought stress could significantly induce *OsbZIP86* expression, and *OsbZIP86* also responded to salt stress (Supplemental Figure **S2C**,**D**). Our qRT-PCR results also showed that expression of *OsbZIP86* was induced, whereas miR2105 expression was repressed after ABA and salt treatments in ZH11 seedlings (Figure **1A,C**). *OsbZIP86* and miR2105 were up- and downregulated, respectively after 0.5– 2 h of water-pouring-out treatment (Figure **1B)**. miR2105 expression rapidly increased within 0.5–1 h recovery; by contrast, *OsbZIP86* expression returned to its normal level after 2–12 h of re-watering (Figure **1B**). Thus, *OsbZIP86* showed the opposite trend of expression change to that of miR2105 in ZH11 seedlings under ABA, salt, and drought treatments. To further evaluate the expression pattern of *OsbZIP86* in different tissues, qRT-PCR was performed. The results showed that *OsbZIP86* was mainly expressed in the stem, followed by the leaf sheath and young leaf (Supplemental Figure **S2B**), similar to the results from the Rice eFP database (Supplemental Figure **S2E**,**F**). GUS staining of *OsbZIP86pro: GUS* rice also showed that *OsbZIP86* was highly expressed in young leaf, leaf sheath, stem, and seed, with lower expression in the panicle (Supplemental Figure **S2G**), consistent with the qRT-PCR data (Supplemental Figure **S2B**). The temporal and spatial expression patterns of miR2105 and *OsbZIP86* in ZH11 also showed highly negative correlations at the same growth phases (Supplemental Figure **S2B**). Therefore, we concluded that ABA, salt, and drought repressed expression of miR2105, resulting in increased expression of *OsbZIP86*.

To test the subcellular localization of OsbZIP86, its the coding sequence (CDS) was fused with *GFP* and introduced into rice protoplasts using PEG (Supplemental Figure **S3A**). The recombinant OsbZIP86-GFP was found to exclusively co-localize with a nuclear marker, NSL-rk-mCherry (Supplemental Figure **S3A**). The stable *Ubi: OsbZIP86-GFP* transgenic rice also showed that the green fluorescence signal of the OsbZIP86-GFP fusion protein was located in the nuclei of root cells (Supplemental Figure **S3B**). These data showed that OsbZIP86 is a nuclear-localized protein.

#### miR2105 and *OsbZIP86* had opposite effects on the tolerance of rice to drought and salt stresses

To determine the detailed functions of *OsbZIP86* and miR2105 in response to different stresses in rice, they were used to generate transgenic rice (Supplemental Figure **S1**), which were subsequently tested with respect to their response to drought and NaCl treatments. Under normal growth conditions (ck), miR2105 and *OsbZIP86* transgenic rice did not show phenotypic difference compared with ZH11, including with respect to plant growth, stomatal area, seed germination, and the main agronomic traits (Figure **2A,D**, **3, 6**, and Supplemental Figure **S4**). However, when these transgenic rice seedlings were recovered after water deficiency stress, the seedling survival rate of ZH11 was about 7%, whereas those of STTM2105 and *bZIP86*-ox seedlings ranged from 15% to 18%, and those of miR2105-ox, *crbzip86*, and *bZIP86*-RNAi seedlings ranged from 0% to 2% (Figure **2A,B**). The leaf water loss rates of STTM2105 and *bZIP86*-ox seedlings were slower whereas those of miR2105-ox, *crbzip86*, and *bZIP86*-RNAi were faster than those of ZH11 under dehydration conditions (Figure **2C**). To test the effects of *OsbZIP86* and miR2105 on agronomic traits under drought conditions, the transgenic rice were grown in boxes filled with sandy with lacunar drought. We found that miR2105-ox, *crbzip86*, and *bZIP86*-RNAi plants showed lower grain weight per panicle, seed setting, and thousand grain weight than ZH11 plants, whereas STTM2105 and *bZIP86*-ox did not show any significant difference compared with ZH11 (Figure **2E–G**). STTM2105 and *OsbZIP86*-ox also enhanced the salt stress tolerance of rice; by contrast, miR2105-ox, *crbzip86*, and *bZIP86*-RNAi decreased salt stress tolerance during the seed germination and seedling stages (Supplemental Figure **S4**). These results indicated that decreased miR2105 expression and increased *OsbZIP86* expression improved the drought and salt tolerance of rice, whereas increased miR2105 expression and decreased *OsbZIP86* expression reduced the drought and salt tolerance of rice.

**Figure 2.**
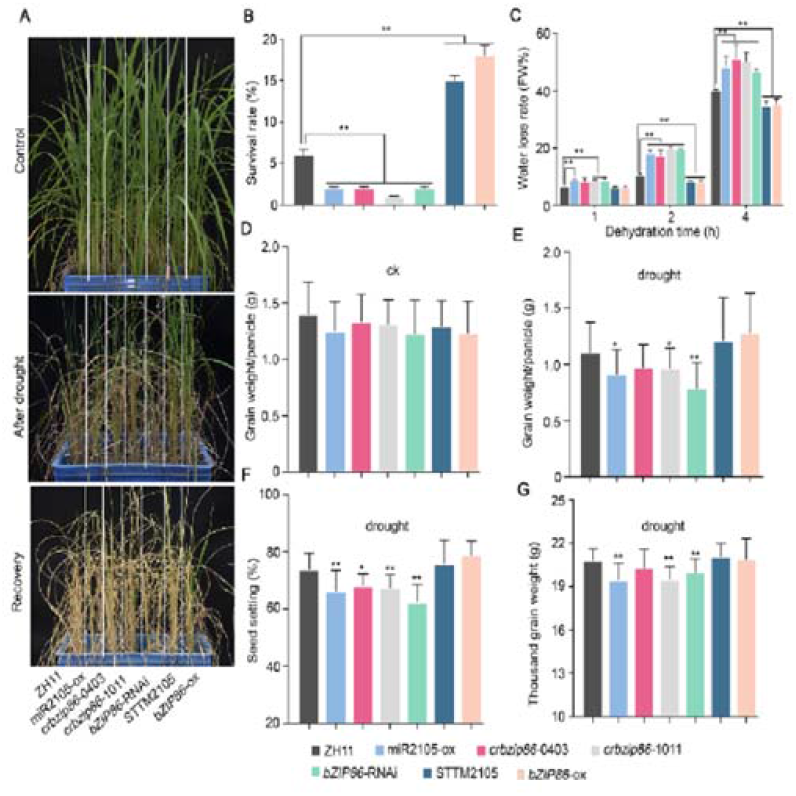
miR2105 and *OsbZIP86* mediate drought-resistance and grain yield of rice under drought condition. (A–B) Phenotypes (A) and survival rates (B) of transgenic rice seedlings under drought treatment. Two-week-old rice seedlings were grown in boxes with sandy soil, water was poured out, and irrigation was stopped for 2 weeks until the leaves wilted for 3 d (middle); then, irrigation was resumed for 1 week (bottom), and the seedlings were watered as the control (top). (C) Water loss rate in detached leaves of the transgenic rice seedlings. Values are means ± SD of 30 independent plants. (D–E) Grain weights per panicle under normal (D) and drought conditions (E). (F–G) Seed setting rate (F) and thousand grain weight (G) under drought treatment. All plants were grown in boxes filled with sandy soil. For drought treatment (F–G), all plants were grown under normal conditions until flowering, then all the water in the boxes was poured out and watering was stopped for 1 week, before plants were recovered with water for 3 d. Lacunar drought treatment was carried out from flowering to mature grain. The experiments were performed in three replicates with similar results, and two independent lines of each transgenic construction were tested. Each repeat was measured in at least 30 seedlings in (A–B) and in 20 independent plants in (D–G). Values are means ± SD; \**p* < 0.05, \*\**p* < 0.01 according to student’s *t-*test in (B–G). miR2105-ox, miR2105 overexpression; *cribzip86-0403*/-1011, *OsbZIP86*-CRISPR; *bZIP86*-RNAi, *OsbZIP86* RNAi; STTM2105, miR2105 downregulation; *bZIP8686*-ox, *OsbZIP86* overexpression.

**Figure 3.**
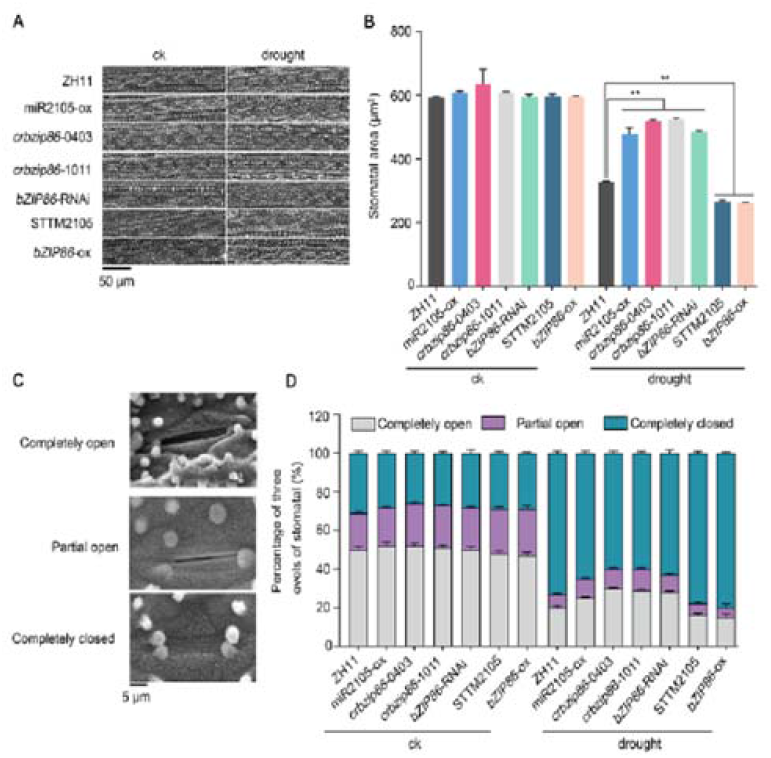
miR2105 and *OsbZIP86* mediate leaf stomatal opening in rice. (A) Stomatal arrangement in abaxial leaf blade. (B) Area per stomatal pore. (C) SEM images of three levels of stomatal opening. (D) Statistics of stomatal opening state. Eight-week-old seedling leaves of ZH11, miR2105, and *OsbZIP86* transgenic rice plants were measured before (ck) and after drought stress. All images were continuously observed by SEM. Six seedlings of each line were used for measurement, and 180 stomata per line were measured (B and D). The experiments were performed in three biological replicates with similar results. Values shown are means ± SD of six independent seedlings. \**p* < 0.05, \*\**p* < 0.01 according to student’s *t-*test.

**Figure 4.**
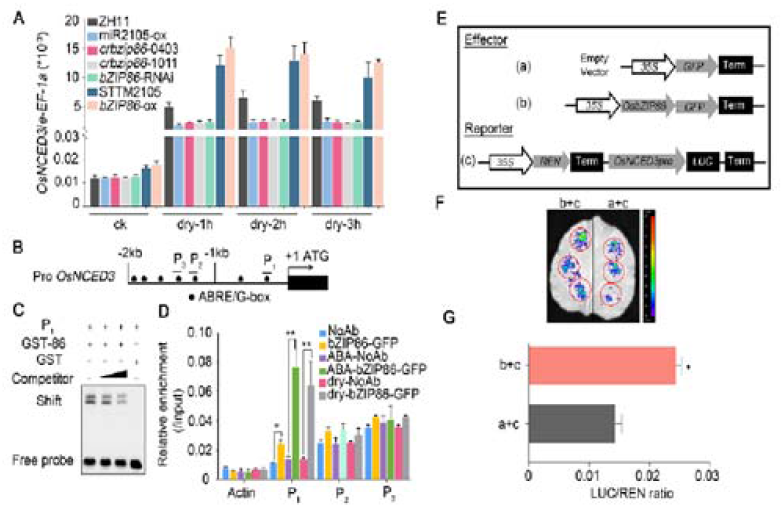
OsbZIP86 binds to the G-box of *OsNCED3* promoter to regulate its expression. (A) Expression levels of *OsNED3* in miR2105 and *OsbZIP86* transgenic rice under normal conditions (ck) and drought stress. The rice *e-EF-1a* gene was used as the internal control. Data represent means ± SD (n = 3). All qRT-PCR analyses for gene expression were performed in three biological replicates with similar results. (B) G-box elements (black dot) in 2-kb *OsNCED3* promoter region. P_1_, P_2_, and P_3_ represent probe positions for EMSA and amplification regions for ChIP-qPCR. (C) *In vitro* EMSA using G-box sequences from promoter of *OsNCED3* as probes. The P_1_ probe was a biotin-labelled fragment of the *OsNCED3* promoter, and the competitor was a non-labelled competitive probe. GST-tagged OsbZIP86 was purified, and 2 μg protein was used. The gradient indicates the increasing amount of competitor. GST-86, fusion protein GST-OsbZIP86; GST, negative control. (D) *In vivo* ChIP-qPCR using ZH11 (NoAb, no antibody) and OsbZIP86-GFP overexpressing line (bZIP86-GFP). The anti-GFP antibody was used to precipitate DNA bound to OsbZIP86. Precipitated DNA was amplified with primers overlapping the G-box motif (P_1_, P_2_, and P_3_). For drought and ABA treatment, ZH11 and *OsbZIP86*-6HA-GFP lines were grown in boxes filled with Yoshida solution for 2 weeks, then water was poured out or they were treated with 50 μm ABA for 2 h. Values are means ± SD from three parallel repeats. NoAb served as a negative control. Rice *Actin* was used as an internal control. (E) Schematic diagram of various constructs for *in vivo* luciferase transient transcriptional activity assay. *35S: OsbZIP86-GFP* was constructed as the effector. *35S: REN*-*OsNCED3pro: LUC* was constructed as the reporter. Free GFP (empty vector) was used as a negative control. (F–G) *In vivo* luciferase activity assay in tobacco leaves. D-luciferin was used as the substrate of luciferase. The expression level of *REN* was used as an internal control. The LUC/REN ratio represents the relative activity of the *OsNCED3* promoter. Error bars indicate SD with biological triplicates (n = 3). \**p* < 0.05, \*\**p* < 0.01, according to student’s *t-*test in (D and G).

**Figure 5.**
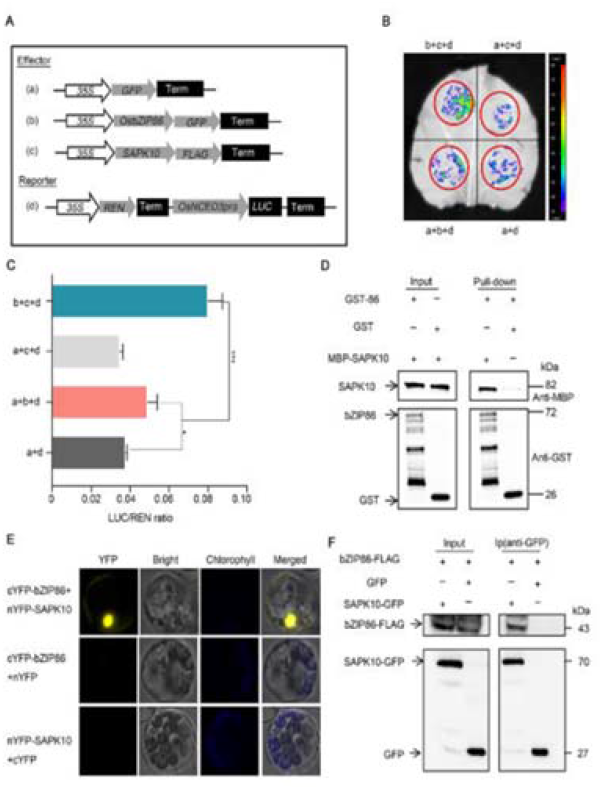
OsbZIP86 interacts with and functions cooperatively with OsSAPK10 to positively regulate *OsNCED3* expression. (A) Schematic diagram of various constructs used in the transient transformation assay. Free GFP was used as a negative control. (B–C) *In vivo* luciferase activity assay in tobacco leaves. D-luciferin was used as the substrate of luciferase. The expression level of *REN* was used as an internal control. The LUC/REN ratio represents the relative activity of the *OsNCED3* promoter. Error bars indicate SD with biological triplicates (n = 3). \**p* < 0.05, \*\*\**p* < 0.001 according to student’s *t-*test. (D) BiFC analysis of interaction between OsbZIP86 and OsSAPK10 *in vivo*. CDS of *OsbZIP86* and *OsSAPK10* fused with the C-terminus and the N-terminus of yellow fluorescent protein (YFP) were co-transformed into rice protoplasts. Overlaid images show signals for YFP (yellow) and chloroplasts (blue). nYFP alone was used as a negative control. (E) Pull-down analysis of interaction between OsbZIP86 and OsSAPK10 *in vitro*. MBP-OsSAPK10 was incubated with GST or GST-OsbZIP86 proteins, and the immunoprecipitated proteins were detected by an anti-GST antibody. Free GST was used as the negative control. (F) Co-IP analysis of interaction between OsbZIP86 and SAPK10 *in vivo*. GFP, OsSAPK10-GFP, and OsbZIP86-FLAG were co-expressed in tobacco leaves by *Agrobacterium* injection. Total protein extracts were immunoprecipitated with the immobilized anti-GFP antibody (Ip), and the immunoprecipitated protein was then detected by using an anti-FLAG antibody. Input OsSAPK10-GFP and OsbZIP86-FLAG proteins were detected with anti-GFP and anti-FLAG antibodies, respectively. The molecular weights (kDa) and proteins are indicated in the left and right panels, respectively.

**Figure 6.**
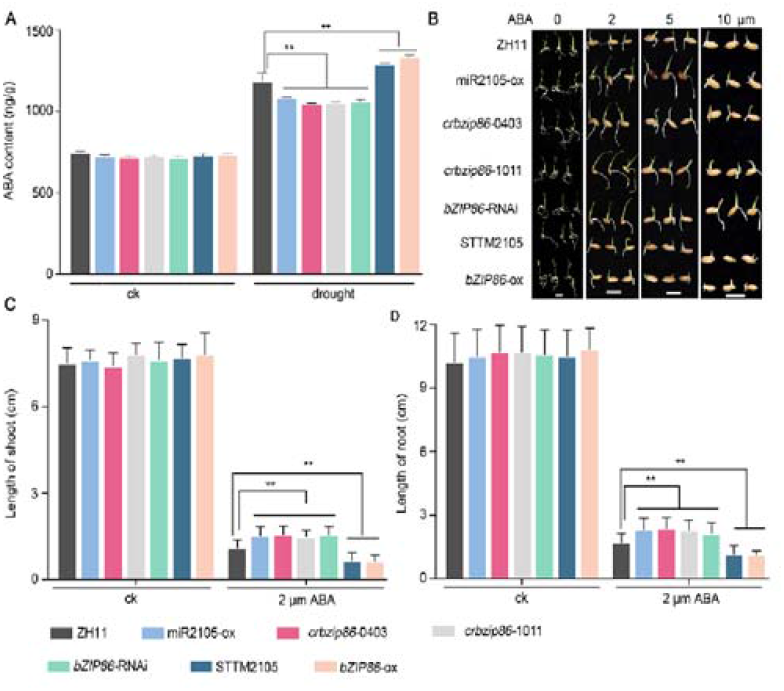
miR2105 and *OsbZIP86* mediate leaf ABA biosynthesis of rice under drought conditions. (A) Leaf ABA content of miR2105 and OsbZIP86 transgenic rice under normal (ck) and drought stress conditions. Error bars indicate SD for biological triplicates (n = 3). (B) Germination performance of miR2105 and OsbZIP86 transgenic rice seedlings under ABA treatment. Seeds were placed on double sheets of filter paper in a 9-cm Petri dish and moistened with distilled water or 2, 5, or 10 μM ABA for 7 d. Scale bar, 0.5 cm. (C–D) Lengths of shoots (C) and main roots (D) of miR2105 and *OsbZIP86* transgenic rice seedlings treated with distilled water or 2 μM ABA for 7 d. Experiments were performed using three biological replicates with similar results. Each repeat was measured in 30 independent seedlings. Values show means ± SD of 30 independent plants. \**p* < 0.05, \*\**p* < 0.01 according to student’s *t-*test (A, C, and D).

When the transgenic rice were grown under water deficiency conditions, the stomatal areas of miR2105-ox, *crbzip86*, and *bZIP86*-RNAi plants were smaller whereas those of STTM2105 and *bZIP86*-ox plants were larger than those of ZH11 (Figure **3A,B**). In addition, more stomata were completely closed and fewer stomata were completely open in STTM2105 and *bZIP86*-ox leaves compared with ZH11. The opposite tendency was observed for miR2105-ox, *crbzip86*, and *bZIP86*-RNAi (Figure **3C,D**). These results indicate that miR2105 and *OsbZIP86* plays positive and negative roles in stomata movement under water deficiency, respectively (Figure **3**).

### OsbZIP86 directly binds to the promoter of *OsNCED3*

Given that OsbZIP86 had been demonstrated to bind to the G-box motif of the wheat *Em* promoter (Nantel & Quatrano, 1996), and promoters of ABA biosynthetic and metabolic genes (Huang *et al*., 2018) including *OsNCED1–*5 and *OsABA8ox1*–*3* harbored G-box *cis*-elements, we checked whether overexpression of *OsbZIP86* could change the expression of these genes. Under normal conditions, none of these genes showed significant differences in expression between ZH11 and *bZIP86*-ox; only *OsNCED3* and *OsABA8ox2* showed slight upregulation in *bZIP86*-ox compared with ZH11 (Supplemental Figure **S5C**,**G**). However, under drought treatment, expression of *OsNCED3* and *OsABA8ox2* was significantly upregulated, especially that of *OsNCED3* (Supplemental Figure **S5C**,**G**). By contrast, the expression of *OsNCED5* was downregulated in *bZIP86*-ox compared with ZH11 (Supplemental Figure **S5E**). To investigate whether miR2105 affected *OsNCED3* expression, *OsNCED3* mRNA levels were further analyzed in leaves of miR2105 transgenic rice under normal and drought stress conditions. Under normal conditions, *bZIP86*-ox and STTM2105 showed slight upregulation of *OsNCED3* compared with ZH11 (Figure **4A**). However, levels of the *OsNCED3* transcript were significantly increased after drought treatment in STM2105 and *bZIP86*-ox seedlings, but significantly decreased in miR2105-ox, *crbzip86*, and *bZIP86*-RNAi seedlings compared with those in ZH11 (Figure **4A**). These results indicate that OsbZIP86 could promote *OsNCED3* transcription under drought conditions.

To further investigate whether OsbZIP86 could directly bind to the promoter of *OsNCED3*, we performed Electrophoretic mobility shift assay (EMSA) and Chromatin immunoprecipitation-quantitative PCR (ChIP-qPCR) to test the DNA-binding ability of OsbZIP86 to the promoter of *OsNCED3*. Based on the G-box motifs present in the *OsNCED3* promoter (Figure **4B**), we designed three pairs of specific primers for ChIP-qPCR and three labeled probes for EMSA (Supplemental Table **S2** and Figure **4B**). We found that OsbZIP86 could specifically bind to the probe 1 (P_1_) region of the *OsNCED3* promoter (Figure **4C** and Supplemental Figure **S6A**). EMSA binding was substantially weakened by non-labeled competitive probe in a dosage-dependent manner (Figure **4C**). Subsequently, ChIP-qPCR was performed to validate this binding *in vivo*, using an anti-GFP antibody and specific primers. Consistent with the EMSA results, OsbZIP86 was enriched in the P_1_ region of the *OsNCED3* promoter (Figure **4D**). Moreover, the enrichment was significantly enhanced under drought conditions and ABA treatment (Figure **4D**). These results showed that OsbZIP86 could specifically bind to the P_1_ fragment of the *OsNCED3* promoter, which is 387 bp upstream of the ATG of the *OsNCED3* coding region (Figure **4B–D** and Supplemental Figure **S6A**). Finally, a dual-luciferase reporting system in tobacco leaves was used to determine the regulatory effect of OsbZIP86 on *OsNCED3* transcription (Figure **4E–G**). In comparison with the empty effector, LUC activity was significantly enhanced when *OsNCED3pro: LUC* was co-transfected with *35S: OsbZIP86* (Figure **4F–G**). Therefore, we concluded that OsbZIP86 could bind to the *OsNCED3* promoter to active its transcription, and it was activated by drought and ABA to promote the transcription of *OsNCED3*.

### OsbZIP86 interacts with OsSAPK10 to positively regulate *OsNCED3* expression

The SAPKs, which are components of ABA signaling, have been reported to activate bZIP TFs in plants (Banerjee & Roychoudhury, 2017; Fu *et al*., 2021). To test whether OsbZIP86 interacted with OsSAPK1-10, luciferase complementation imaging (LCI) in tobacco was performed (Supplemental Figure **S6B**). We found that OsbZIP86 could interact with OsSAPK4, OsSAPK6, OsSAPK7, and OsSAPK10. Then, we performed a dual-luciferase transient transcriptional activity assay to examine whether the four OsSAPKs could function cooperatively with OsbZIP86 to positively regulate *OsNCED3* expression. We detected a significant increase in *OsNCED3* promoter activity when *OsSAPK4, OsSAPK7*, and *OsSAPK10* were co-transformed with *OsbZIP86* (Figure **5A–C** and Supplemental Figure **S6C**,**D**). The transcription level of the *OsNCED3* promoter was most strongly increased by OsSAPK10 (Figure **5A–C**). The interaction between OsbZIP86 and OsSAPK10 was further confirmed by a pull-down assay *in vitro* and by bimolecular fluorescence complementation (BiFC) and Co-immunoprecipitation (Co-IP) assays *in vivo* (Figure **5D–F**). These results indicate that OsSAPK10 may activate OsbZIP86 and thus cooperatively regulate *OsNCED3* expression. OsSAPK10 has been reported to positively regulate ABA sensitivity (Min *et al*., 2007; Wang *et al*., 2020). Together, we suggested that OsSAPK10 facilitates OsbZIP86 to positively regulate the transcript expression of *OsNCED3*.

### miR2105 and *OsbZIP86* mediate ABA biosynthesis

To test whether endogenous ABA biosynthesis was mediated by miR2105 and *OsbZIP86*, we measured the ABA contents of leaves of transgenic rice (Figure **6A**). When the rice seedlings were grown under normal conditions, the ABA contents of transgenic rice did not show any difference compared with those of ZH11 rice (Figure **6A**). However, when they were treated by drought for 4 h, the ABA contents of miR2105-ox, *crbzip86*, and *bZIP86*-RNAi plants were lower compared with ZH11, whereas those of STTM2105 and *bZIP86*-ox were higher (Figure **6A**). To investigate whether exogenous ABA treatment could complement the change in ABA biosynthesis in these transgenic rice, we performed ABA treatment (Figure **6B–D**). The seedling growth of these transgenic rice plants did not show any difference compared with that of ZH11 under normal conditions (Figure **6B–D**).

However, after ABA treatment, the seedlings of miR2105-ox, *crbzip86*, and *bZIP86*-RNAi plants had longer shoots and roots, and those of STTM2105 and *bZIP86*-ox plants had shorter shoots and roots compared with ZH11 (Figure **6B–D**). This may indicate that exogenous ABA treatment strongly repressed ZH11 growth and slightly complemented the endogenous ABA insufficiency of miR2105-ox, *crbzip86*, and *bZIP86*-RNAi plants. Conversely, there was too much ABA in STTM2105 and *bZIP86*-ox plants after ABA treatment, and this further repressed the growth of these seedlings. Therefore, we propose that miR2105 and *OsbZIP86* regulate ABA biosynthesis only in the presence of drought stress.

## Discussion

Plants quickly accumulate ABA to activate a series of stress responses when subjected to abiotic stresses. When environmental conditions are optimal, the amount of ABA is reduced to basic levels that promote optimal growth. When plants encounter a non-optimal environment, the regulation of ABA level in tissues and cells is essential for balancing defense and growth (Chen *et al*., 2020). Therefore, excessive ABA levels under normal conditions will adversely affect the normal growth of plants. Here, we found that miR2105 targeted *OsbZIP86*, regulating its expression at post-transcriptional levels. OsbZIP86 directly activates *OsNCED3* transcription by binding to the G-box in the promoter, and interacts with OsSAPK10, resulting in enhanced-*OsNCED3* expression to control ABA biosynthesis. Therefore, the drought tolerance of the stable miR2105-overexpressing or *OsbZIP86*-downregulated transgenic rice plants was enhanced without any penalty with respect to major agronomic traits under normal conditions. Our findings suggest a molecular breeding strategy for improving the drought resistance of rice without affecting agronomic traits by using miR2105 or *OsbZIP86*.

### miR2105/*OsbZIP86* regulates drought-induced ABA biosynthesis through *OsNCED3*

Under water stress, the ABA content in rice leaves can be rapidly induced within 1 h and quickly decrease to the basal line within 1 h during rehydration (Ye *et al*., 2011). NCEDs have been reported to be involved in the rate-limiting step in the ABA biosynthetic pathway (Nambara & Marion-Poll, 2005; Chen *et al*., 2020). However, how to regulate expression of *NCEDs* only under abiotic stresses was not completely clear. Our data showed that the expression of miR2105 in ZH11 plants was repressed by drought, ABA, and salt treatments; conversely, the expression of *OsbZIP86* was induced (Figure **1A–C**). We found that miR2105 could repress the expression of *OsbZIP86*, and miR2105 could direct to cleave *OsbZIP86* mRNA (Figure **1D,E**). In addition, the transgenic rice plants with changed expression of *OsbZIP86* and miR2105 showed the expected phenotypic changes under drought conditions (Figure **2**, **3** and **6**): miR2105 negatively regulated the drought/salt tolerance and ABA sensitivity of the seedlings, whereas *OsbZIP86* positively regulated these responses (Figure **2**, **6** and Supplemental Figure **S4**). Therefore, we concluded that *OsbZIP86* is a target gene of miR2105 and its expression level is regulated by miR2105 under drought salt and ABA treatment.

Rice is predicted to have five *OsNCED*s (Huang *et al*., 2018; Hwang *et al*., 2018; Huang *et al*., 2019), which are tightly regulated in response to developmental or stress conditions (Chen *et al*., 2020). *OsNCED3*, which encodes a key enzyme in ABA synthesis, has the G-box core sequence 5′-ACGT-3′ in its promoter, and its expression is significantly induced by NaCl, PEG, and H_2_O_2_ stresses (Huang *et al*., 2018). *OsbZIP86* is predicted to be a G-box binding factor, and it can bind to and activate the wheat *Em* promoter (Nantel & Quatrano, 1996). Our EMSA, ChIP-qPCR, and luciferase activity assays showed that *OsbZIP86* could directly bind to the *OsNCED3* promoter (Figure **4**). We also demonstrated that drought and ABA treatment could enhance the binding ability of *OsbZIP86* to the *OsNCED3* promoter (Figure **4D**), suggesting that the function of OsbZIP86 might be dependent on the ABA signaling pathway. However, the stable transgenic rice plants with altered expression of *OsbZIP86* and miR2105 showed significant differences in *OsNCED3* expression (Figure **4A**), ABA content (Figure **6A**), and drought tolerance (Figure **2**) only when the plants were treated by drought. Of the five rice *OsNCEDs*, only *OsNCED3* showed significant upregulation of expression in *OsbZIP86*-ox plants under drought conditions (Supplemental Figure **S5**). Therefore, we concluded that *OsbZIP86* is a direct regulator of *OsNCED3*, and that *OsNCED3* expression is regulated under drought stress.

Like *OsNCED3, OsNCED5* is expressed in all tissues and regulates ABA biosynthesis and tolerance of rice to salt and water stresses (Huang *et al*., 2019). However, overexpression of *OsbZIP86* could cause upregulation of *OsABA8ox2* and downregulation of *OsNCED5* under drought treatment (Supplemental Figure **S5**). *OsbZIP86* has ABRE *cis*-elements for ABA signaling (Figure **1F**), suggesting that *OsbZIP86* may positively regulate ABA signaling. The strongly induced expression of *OsNCED3* in *OsbZIP86*-ox (Supplemental Figure **S5C**) under drought conditions in turn may negatively regulate the expression of *OsNCED5* and positively regulate expression of *OsABA8ox2* to maintain the balance of endogenous ABA.

### OsSAK10 interacts with OsbZIP86 to regulate the transcript expression of *OsNCED3*

SnRK2 kinases have been reported to play essential parts in ABA signaling through phosphorylating AREB/ABF TFs at the post-translational level in *Arabidopsis* (Banerjee & Roychoudhury, 2017). Several OsSAPKs have been found to phosphorylate rice bZIPs via protein–protein interactions, for instance, OsSAPK6 can interact with OsbZIP10 (Chae *et al*., 2007) and OsbZIP46 (Tang *et al*., 2012; Kim *et al*., 2015), OsSAPK2 can interact with OsbZIP23 and OsbZIP46 (Zong *et al*., 2016), OsbZIP72 can interact with OsSAPK10 and OsSAPK1 (Wang *et al*., 2020; Fu *et al*., 2021). We found that OsbZIP86 interacted with OsSAPK10 (Figure **5D–F** and Supplemental Figure **S6B**). Overexpression of *OsSAPK10* reduced water loss in detached leaves (Min *et al*., 2019) and increased sensitivity to ABA (Wang *et al*., 2020). These phenotypes were similar to those resulting from overexpression of *OsbZIP86* (Figure **2** and **6**). Moreover, the expression pattern and subcellular localization of OsSAPK10 (Wang *et al*., 2020) overlapped with those of OsbZIP86 (Supplemental Figure **S2B** and Supplemental Figure **S3**). OsSAPK10 could function cooperatively with OsbZIP86 to positively regulate *OsNCED3* expression (Figure **5A–C**). These results indicate that OsbZIP86 is activated by OsSAPK10. Taken together, our results suggest that OsbZIP86 may regulate ABA biosynthesis through *OsNCED3* in an OsSAPK10-dependent manner. Further research is needed to investigate how OsSAK10 enhances the activity of OsbZIP86.

### Overexpression of *OsbZIP86* and downregulation of miR2105 can improve drought tolerance without affecting the main agronomic traits of rice

ABA can improve plant tolerance to abiotic stresses, but ABA induces leaf senescence and negatively affects the yield of rice. Direct overexpression of ABA biosynthetic genes (such as *OsNCED3*) or its biosynthetic regulators (such as *OsNAC2*) caused negative effects on leaves and yield, while these transgenic plants could improve the stress tolerance (Mao *et al*., 2017; Huang *et al*., 2018). Therefore, when the ABA synthesis is promoted only under stress (such as drought), the resistance of plants can be improved without affecting their growth and development under normal conditions. Thus, these genes could be used in molecular breeding to improve stress tolerance without negative effects.

The transgenic rice with altered expression of miR2105 or *OsbZIP86* showed changes in ABA contents associated with overexpression of *OsNCED3* only when drought was present (Figure **6A**), leading to changes in the drought resistance in rice seedlings (Figure **2** and **3**). These data indicate that miR2105 and *OsbZIP86* regulate ABA biosynthesis to enhance drought tolerance through *OsNCED3* only under drought conditions, while not affecting aspects of rice growth including plant height, tiller numbers, seed setting percentage, and thousand seed weight under normal conditions (Supplemental Figure **S7**). However when treated with lacunar drought and salt, the seed setting percentage, thousand seed weight, and grain yield per panicle of *crbzip86, bZIP86*-RNAi, and miR2105-ox were lower than those of ZH11 (Figure **2E–G** and Supplemental Figure **S4G–I**). miR2105-ox and *bZIP86*-ox showed no significant differences compared with ZH11, possibly owing to the insufficient drought and salt treatments. These results imply that altered expression of miR2105 or *OsbZIP86* had no impact on rice growth and development under normal conditions but could improve the stress resistance of rice under drought and salt stresses; this is different from other genes for they may exert negative effects on normal rice growth when promoting ABA biosynthesis (Kasuga *et al*., 2004; Ito *et al*., 2006). Therefore, miR2105 and OsbZIP86 may have potential for use in promoting drought tolerance without penalty to major agronomic traits under normal conditions.

In summary, we have proposed a working model of how OsbZIP86 is regulated by miR2105 and OsSAK10 to control ABA biosynthesis through *OsNCED3* and functions in regulating drought resistance in rice (Figure **7**). Under drought stress, the expression of miR2105 is repressed, whereas the expression of *OsbZIP86* is induced. Then, OsbZIP86 is activated by OsSAPK10 through direct interaction, thereby promoting the expression of *OsNCED3* by directly binding to the promoter. The upregulated *OsNCED3* plays an important part in the regulation of ABA biosynthesis to modulate drought resistance.

**Figure 7.**
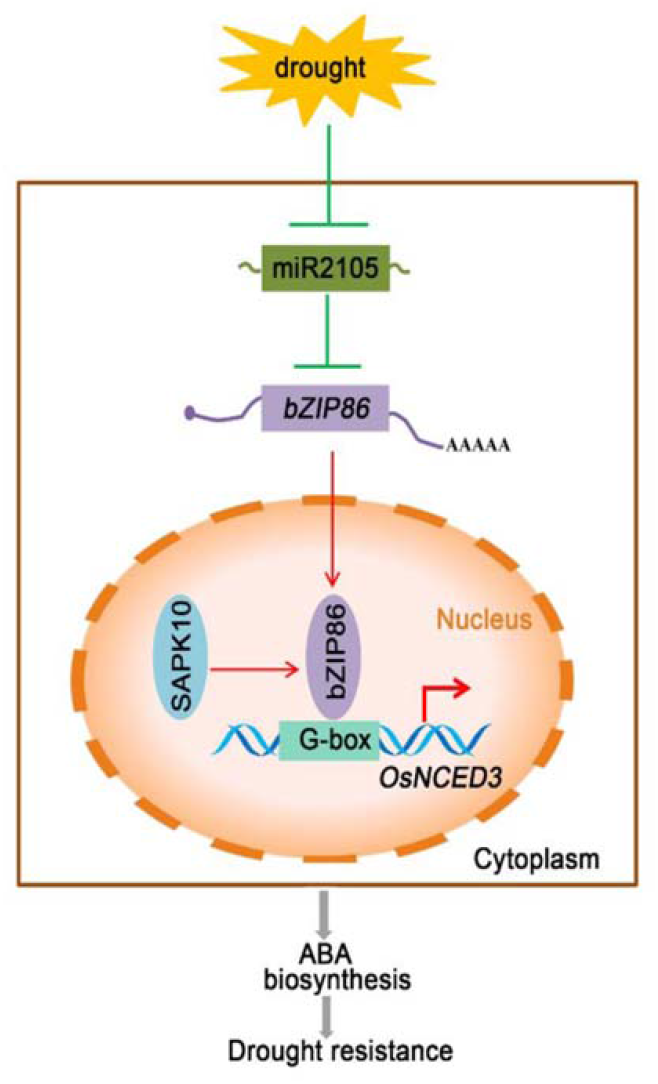
Working model of ‘miR2105-OsbZIP86-*OsNCED3*’ module functions in regulating drought resistance Under drought stress, the expression of miR2105 is repressed, whereas the expression of *OsbZIP86* is induced. Then, OsbZIP86 is activated by OsSAPK10 through direct interaction, thereby promoting the expression of *OsNCED3*. The upregulated *OsNCED3* has an important role in regulating ABA biosynthesis to modulate drought resistance. Arrowheads show positive regulation; flat-ended lines show negative regulation.

## Materials and Methods

### Vector construction and rice transformation

To generate osa-miR2105-overexpression rice (miR2105-ox), a mature 20-base-pair (bp) sequence (MI0010566) of miR2105 was downloaded from miRBase (http://www.mirbase.org). The cloning procedure for miR2105-ox followed the description at http://wmd3.weigelworld.org/cgi-bin/webapp.cgi. The artificial osa-miR2105 was inserted downstream of the *35S* promoter in pCAMBIA1301 (http://www.cambia.org). To downregulate osa-miR2105 (STTM2105), the short tandem target mimic method was used as described previously (Yan *et al*., 2012). The fragment (GAAGCTTT<u>TTGTGATGTGAATGATTCAT</u>GTTGTTGTTGTTATGGTCTAGTTGTTGTTGTTATGGTCTAATTTAAATATGGTCTAAAGAAGAAGAATATGGTCTAAAGAAGAAGAAT<u>TTGTGATGTGAATGATTCAT</u>GGATCCA) was synthetized by Invitrogen™ (China) and inserted downstream of the *Ubi-1* promoter in pXU1301 (modified from pCAMBIA1301; the *35S* promoter was replaced by the *Ubi-1* promoter). Therefore, the fragment contained two target mimic sequences (underlined).

For overexpression of *OsbZIP86* (*bZIP86*-ox), the CDS of *OsbZIP86* was amplified from *Oryza sativa* L. cv “Zhonghua 11” (ZH11) and subcloned into pXU1301 to produce a 6×HA–GFP fusion protein expressed from the *Ubi* promoter. To generate *OsbZIP86*-RNAi, two fragments of the *OsbZIP86* CDS (391 bp) were inserted downstream of the *Ubi-1* promoter in the rice RNA interference (RNAi) vector pTCK303 (Wang *et al*., 2004). The *osbzip86* mutants (cr*bzip86*) were generated by a CRISPR/Cas9 genome-editing system (Ma *et al*., 2015). The targets selected are listed in Supplemental Figure **S1E**. Individual T_0_ transformants were analyzed by sequencing their *OsbZIP86* target regions, which were amplified by PCR. To generate *OsbZIP86pro: GUS* transgenic plants, an approximately 2-kb promoter sequence of *OsbZIP86* was cloned and inserted into pCAMBIA1301.

All the above-mentioned constructs were introduced into *Agrobacterium tumefaciens* strain *EHA*105, and ZH11 was transformed by *Agrobacterium*-mediated transformation. All the primers used in this study are listed in Supplemental Table **S2**.

### 5′-RLM-RACE assay

Total RNA from tillering stage leaf sheaths of ZH11 was directly ligated to a synthesized RNA adaptor (GCUGAUGGCGAUGAAUGAACACUGCGUUUGCUGGCUUUGAUGAAA). The cDNA was amplified by nested PCR, and the final PCR product was gel-purified and subcloned into the pGEM-T Easy Vector (Promega, Guangzhou, China) for sequencing. Other processes for 5′-RLM-RACE were as previously described (Xia *et al*., 2015a). The primers are listed in Supplemental Table **S2**.

### Plant growth conditions and stress treatments

All the rice used in this study was generated from ZH11. The seeds were surface-sterilized in 70% (v/v) ethanol for 30 s and then in 2.5% NaClO (w/v) solution for another 40 min, followed by several rinses with sterile water. Then, seeds were incubated in darkness at 28 °C for 2 d to induce germination. The rice were grown under controlled field conditions in Guangzhou, China, or in boxes filled with Yoshida solution at 30 °C with a 14 h/10 h light/dark period.

To determine the effects on gene expression of ABA, water deficiency, and salt treatments, 2-week-old ZH11 seedlings grown in Yoshida solution were supplied with 0.5 μM ABA for different times, exposed to air for 4 h, or treated with 150 mM NaCl for 4 h. The seedlings were then transferred to Yoshida solution again for recovery.

For drought tolerance test, 2-week-old rice seedlings were grown in boxes filled with sandy soil and grown for 2 weeks; then, all the water in the boxes was poured out and watering was stopped for 2 weeks until the seedling wilted for 3 d and recovery 1 week. Photographs were taken and the surviving seedlings were counted.

For the ABA sensitivity test at the seedling stage, transgenic rice seeds were placed on double sheets of filter paper in a 9-cm Petri dish and moistened with distilled water or different concentrations of ABA (2, 5, or 10 μM) for 7 d. Lengths of shoots and roots were measured. Three replicates were tested for each plant line. Thirty seeds were measured for each replicate. Leaf ABA contents were measured using a Phytodetek competitive ELISA kit (Mlbio, Shanghai, China). ABA was extracted as described previously (Fukumoto *et al*., 2013).

To examine the effects of salt on seedling growth, 4-week-old-seedlings were transferred to Yoshida solution with 150 mM NaCl and allowed to grow for 10 d. Photographs were taken and the surviving seedlings were counted.

### Water loss rate and stomatal closure

To detect the water loss rate under dehydration conditions, leaves of 2-week-old rice seedlings were cut into 1-cm lengths, exposed to air at room temperature in the dark, and weighed at different times; the water loss rate was calculated as the percentage of total weight lost compared with the initial weight at each time point (Miao *et al*., 2020). Stomatal closure was examined according to the method described previously (Miao *et al*., 2020). Leaves from 8-week-old rice grown under normal or drought stress conditions were cut in 0.1-cm^2^ pieces and fixed in 2.5% glutaraldehyde solution with 0.1 M phosphate-buffered saline (pH 7.2). Stomatal status was monitored by scanning electron microscopy (SEM) (SU8010, Hitachi, Japan); the SEM analysis was performed as described previously (Xia *et al*., 2015b). Six individual plants from the ZH11 and transgenic rice lines were used for measurements of the stomatal aperture, with 30 stomata measured per plant.

### RNA extraction and reverse transcription PCR (RT-PCR)

The extraction of small RNA and total RNA from rice, reverse transcription and quantitative RT-PCR (qRT-PCR) amplification were performed as previously described (Xia *et al*., 2015a). *OsbZIP86* expression during the plants’ entire growth life was verified using data from RiceXpro (http://ricexpro.dna.affrc.go.jp/).

### β-glucuronidase (GUS) assay

The GUS assay was carried out as described previously (Liu *et al*., 2021) with minor modifications. Transgenic plant tissues were incubated in GUS staining solution (Sangon, Shanghai, China). Following vacuum infiltration, the samples were incubated at 37 °C for 16 h. After staining, the tissues were bleached with 70% ethanol and photographed under a light microscope (Leica DVM6, Leica Microsystems, Germany).

### Subcellular localization of OsbZIP86

To determine the subcellular localization of OsbZIP86, the CDS of *OsbZIP86* was cloned into the pUC18 vector, which we had modified to produce an OsbZIP86-GFP fusion construct driven by the CaMV*35S* promoter (*35S: OsbZIP86-GFP*). Rice protoplasts prepared from etiolated shoots were co-transformed with *35S: OsbZIP86-GFP* and *NSL-mCherry* (a nuclear marker) using polyethylene glycol (PEG). In addition, the CDS of *OsbZIP86-GFP* was cloned into the pXU1301 vector, and the fusion construct (*Ubi: OsbZIP86-GFP*) was introduced into *EHA105* to create transgenic rice. The fluorescence signal was observed with a confocal laser scanning microscope (Leica SP8 STED 3X, Leica Microsystems, Germany). The primers are listed in Supplemental Table **S2**.

### Dual-luciferase assay of transiently transformed tobacco leaves

To generate the luciferase reporter construct for the dual-luciferase assay, the 2-kb promoter sequence of *OsNCED3* was amplified from rice and inserted into a pGreenII0800-LUC vector (Hellens *et al*., 2005) using the *Hind*III and *BamH*I sites, respectively. The CDS of *OsbZIP86* and *OsSAPKs* were cloned into the pCAMBIA1300-GFP and pHB-3×FLAG vectors, respectively, as the effectors. The tobacco leaves were used for dual-luciferase assays as described previously (Hellens *et al*., 2005). The effector construct (*35S: OsbZIP86, 35S: OsSAPKs* or *35S: GFP*) and the reporter construct (*35S: REN*-*OsNCED3pro: LUC*) were introduced into *A. tumefaciens* strain GV3101 and then infiltrated into 3-week-old tobacco leaves by *Agrobacterium* injection. The activities of firefly luciferase (LUC) and Renilla luciferase (REN) were measured using a Dual-Luciferase Reporter Assay System (E2920, Promega, USA). The LUC activity was normalized to the REN activity. The primers are listed in Supplemental Table **S2**.

### EMSA and pull-down assay

The EMSA and pull-down assays were performed as described previously (Wang *et al*., 2020). The CDS of *OsbZIP86* and *OsSAPK10* were cloned into the pGEX4T-1 and pMAL-c2x vectors to generate GST-OsbZIP86 and MBP-OsSAPK10 fusion proteins, respectively. These recombinant vectors were transformed into *Escherichia coli* strain BL21. Fusion proteins were induced with 0.5 mM Isopropyl β-D-Thiogalactoside (IPTG) at 20°C for 12 h. The fusion proteins (GST-OsbZIP86 and MBP-OsSAPK10) were purified using Glutathione Sepharose 4B (GE Healthcare, USA) and Amylose Magnetic Beads (New England Biolabs, USA) according to the manufacturer’s protocols, respectively. For the EMSA assay, complementary single-stranded oligonucleotides derived from 40 bp of the G-box region of the *OsNECD3* promoter were synthesized as DNA probes. The probes were synthesized by Invitrogen (Shanghai, China). EMSA was performed using a LightShift(tm) Chemiluminescent EMSA Kit (Thermo, No. 20148) following the manufacturer’s protocol. For the pull-down assay, GST-OsbZIP86 and MBP-OsSAPK10 were incubated with Glutathione Sepharose 4B (GE Healthcare, USA) in pull-down buffer (50 mM Tris-HCl, pH 7.5, 100 mM NaCl, 0.25% Triton X-100, 35 mM β-mercaptoethanol) at 4 °C for 3 h. After washing six times with pull-down buffer, the pulled proteins were eluted by boiling and further analyzed by immunoblotting using anti-GST. The primers and EMSA probes are listed in Supplemental Table **S2**.

### ChIP-qPCR assay

The ChIP-qPCR assay was performed as described previously (Wang *et al*., 2020). Briefly, 3 g of leaves from 2-week-old seedlings of ZH11 and *OsbZIP86*-6HA-GFP lines grown under normal conditions or treated with 50 μM ABA or subjected to water deficiency for 2 h were cross-linked twice by 3% formaldehyde for chromatin isolation under vacuum for 15 min and stopped using 2 M glycine. Then the samples were ground to powder in liquid nitrogen prior to isolating chromatin. After sonication, the chromatin complexes were incubated with GFP-Trap Agarose (Chromotek, USA). DNA was purified using a PCR purification kit (DNA Clean & Concentrator-5, Tianmo Biotech) and recovered in water for qPCR analysis. Primers were selected in the promoter regions of *OsNCED3* (Supplemental Table **S2**). The amount of precipitated DNA was calculated relative to the total input chromatin and expressed as a percentage of the total according to the formula: % input = 2^ΔCt^ × 100%, where ΔCt = Ct (input) – Ct (IP), and Ct is the mean threshold cycle of the corresponding PCR reaction. The ChIP experiments were repeated three times with the similar results.

### LCI assay

The CDS of *OsbZIP86* and *OsSAPKs* were cloned into the pCAMBIA1300-NLuc and pCAMBIA1300-CLuc LUC vectors (Chen *et al*., 2008), respectively. All of the constructs were introduced into *A. tumefaciens* strain GV3101, which was then co-transformed into 3-week-old tobacco leaves by *Agrobacterium* injection. One millimolar luciferin was sprayed onto leaves to detect the fluorescence signal after 2.5 d infiltration under a plant *in vivo* imaging system (LB985 NightSHADE). The details were examined according to the method described previously (Chen *et al*., 2008).

### BiFC assay

The CDS of *OsbZIP86* and *OsSAPK10* were cloned into the pSATN-cYFP-C1 and pSATN-nYFP-C1 vectors (Citovsky *et al*., 2006), respectively. The constructed vectors were transiently cotransformed into rice protoplasts using PEG. Transfected cells were imaged using a confocal spectral microscope imaging system (Leica, SP8 STED 3X, Germany).

### Co-IP assay

Co-IP was performed as described previously (Wang *et al*., 2020). To generate epitope-tagged expression vectors for Co-IP, the CDS of *OsbZIP86* and *OsSAPK10* were fused with sequences encoding a FLAG tag (vector:pHB-3×FLAG) and GFP tag (vector:pCAMBIA1300-GFP) driven by the *35S* promoter, respectively. The fusion proteins OsbZIP86-FLAG (bZIP86-FLAG) and OsSAPK10-GFP (SAPK10-GFP) were co-transformed into 3-week old tobacco leaves by *Agrobacterium* injection. Total protein was extracted 2.5 d after infiltration with extraction buffer (50 mM Tris-HCl, 150 mM NaCl, 2 mM MgCl_2_, 1 mM DTT, 20% glycerol, 1% NP-40, 0.2 mM PMSF, with complete mini tablet) and then immunoprecipitated with GFP-Trap Agarose (Chromotek, USA) according to the manufacturer’s instructions. The immunoprecipitated proteins were separated via sodium dodecyl sulfate polyacrylamide gel electrophoresis (SDS-PAGE, 10% gel) and analyzed by immunoblotting analysis with anti-GFP or anti-FLAG.

### Statistical analysis

All results were presented as mean ± standard deviation (SD) of three replicates. Statistical analysis was performed using SPSS 19.0, including one-way ANOVA and the least significant difference. Difference was considered significant at *p* < 0.05 and highly significant as *p* < 0.01. Diagrams were prepared using GraphPad Prism 8.3.0 and Adobe Photoshop.

## Supplemental Data

**Supplemental Table S1**. The description of the predicted 13 target genes of miR2105.

**Supplemental Table S2**. Primers used in this study.

**Supplemental Figure S1**. Identification of miR2105 and *OsbZIP86* transgenic rice.

**Supplemental Figure S2**. *OsbZIP86* is a target gene of miR2105 and the expression pattern analyse of *OsbZIP86*.

**Supplemental Figure S3**. Subcellular localization of OsbZIP86.

**Supplemental Figure S4**. miR2105 and *OsbZIP86* mediate salt resistance and grain yield of rice under salt treatment.

**Supplemental Figure S5**. Expression changes of ABA biosynthetic and metabolic genes in *OsbZIP86* overexpression transgenic rice.

**Supplemental Figure S6**. OsbZIP86 binds to promoter fragments of *OsNCED3*, and OsbZIP86 interacts with OsSAPKs to regulate *OsNCED3* expression.

**Supplemental Figure S7**. Agronomic traits of miR2105 and *OsbZIP86* transgenic rice under normal conditions.

## Acknowledgements

We thank Professors Xuncheng Liu (South China Botanical Garden) and Wangjin Lu (South China Agricultural University) for supporting with the *in vitro* dephosphorylation assays. This research was supported by National Natural Science Foundation of China (31971816, 31772384, 32171933); Strategic Priority Research Program of the Chinese Academy of Sciences (XDA24030201-3).

## Parsed Citations

Banerjee A, Roychoudhury A. (2017) Abscisic-acid-dependent basic leucine zipper (bZIP) transcription factors in plant abiotic stress. Protoplasma 254: 3–16.

Bartel DP. (2004) MicroRNAs: Genomics, biogenesis, mechanism, and function. Cell 116: 281–297.

Chae MJ, Lee JS, Nam MH, Cho K, Hong JY, Yi SA, Suh SC, Yoon IS. (2007) Arice dehydration-inducible SNF1-related protein kinase 2 phosphorylates an abscisic acid responsive element-binding factor and associates with ABA signaling. Plant Molecular Biology 63: 151–169.

Chen H, Zou Y, Shang Y, Lin H, Wang Y, Cai R, Tang X, Zhou JM. (2008) Firefly luciferase complementation imaging assay for protein-protein interactions in plants. Plant Physiology 146: 368–376.

Chen K, Li GJ, Bressan RA, Song CP, Zhu JK, Zhao Y. (2020) Abscisic acid dynamics, signaling, and functions in plants. Journal of Integrative Plant Biology 62: 25–54.

Citovsky V, Lee LY, Vyas S, Glick E, Chen MH, Vainstein A, Gafni Y, Gelvin SB, Tzfira T. (2006) Subcellular localization of interacting proteins by bimolecular fluorescence complementation in planta. Journal of Molecular Biology 362: 1120–1131.

Dong T, Park Y, Hwang I. (2015) Abscisic acid: biosynthesis, inactivation, homoeostasis and signalling. Plant Hormone Signalling 58: 29–48.

Finkelstein RR, Gampala S, Rock CD. (2002) Abscisic acid signaling in seeds and seedlings. Plant Cell 14: S15–45.

Fu X, Liu C, Li Y, Liao S, Cheng H, Tu Y, Zhu X, Chen K, He Y, Wang G. (2021) The coordination of OsbZIP72 and OsMYBS2 with reverse roles regulates the transcription of OsPsbS1 in rice. New Phytologist 229: 370–387.

Fukumoto T, Kano A, Ohtani K, Inoue M, Yoshihara A, Izumori K, Tajima S, Shigematsu Y, Tanaka K, Ohkouchi T, et al. (2013) Phosphorylation of D-allose by hexokinase involved in regulation of OsABF1 expression for growth inhibition in Oryza sativa L. Planta 237: 1379–1391.

Hellens RP, Allan AC, Friel EN, Bolitho K, Grafton K, Templeton MD, Karunairetnam S, Gleave AP, Laing WA. (2005) Transient expression vectors for functional genomics, quantification of promoter activity and RNAsilencing in plants. Plant Methods 1: 13.

Huang Y, Guo Y, Liu Y, Zhang F, Wang Z, Wang H, Wang F, Li D, Mao D, Luan S, et al. (2018) 9-cis-epoxycarotenoid dioxygenase 3 regulates plant growth and enhances multi-abiotic stress tolerance in rice. Frontiers in Plant Science 9: 162.

Huang Y, Jiao Y, Xie N, Guo Y, Zhang F, Xiang Z, Wang R, Wang F, Gao Q, Tian L, et al. (2019) OsNCED5, a 9-cis-epoxycarotenoid dioxygenase gene, regulates salt and water stress tolerance and leaf senescence in rice. Plant Science 287: 110188.

Hwang SG, Lee CY, Tseng CS. (2018) Heterologous expression of rice 9-cis-epoxycarotenoid dioxygenase 4 (OsNCED4) in Arabidopsis confers sugar oversensitivity and drought tolerance. Botanical Studies 59: 2.

Ito Y, Katsura K, Maruyama K, Taji T, Yamaguchi-Shinozaki K. (2006) Functional analysis of rice DREB1/CBF-type Transcription factors involved in cold-responsive gene expression in transgenic rice. Plant & Cell Physiology 47: 141–153.

Joo H, Baek W, Lim CW, Lee SC. (2021) Post-translational modifications of bZIP transcription factors in abscisic acid signaling and drought responses. Current Genomics 22: 4–15.

Joo H, Lim CW, Lee SC. (2020) The pepper RING-type E3 ligase, CaATIR1, positively regulates abscisic acid signalling and drought response by modulating the stability of CaATBZ1. Plant, Cell & Environment 43: 1911–1924.

Kasuga M, Miura S, Shinozaki K, Yamaguchi-Shinozaki K. (2004) Acombination of the Arabidopsis DREB1Agene and stress-inducible rd29Apromoter improved drought- and low-temperature stress tolerance in tobacco by gene transfer. Plant & Cell Physiology 45: 346–350.

Kim N, Moon SJ, Min MK, Choi EH, Kim JA, Koh EY, Yoon I, Byun MO, Yoo SD, Kim BG. (2015) Functional characterization and reconstitution of ABA signaling components using transient gene expression in rice protoplasts. Frontiers in Plant Science 6: 614.

Kobayashi Y, Yamamoto S, Minami H, Kagaya Y, Hattori T. (2004) Differential activation of the rice sucrose nonfermenting1-related protein kinase2 family by hyperosmotic stress and abscisic acid. Plant Cell 16: 1163–1177.

Koyama T, Sato F, Ohme-Takagi M. (2017) Roles of miR319 and TCP transcription factors in leaf development. Plant Physiology 175: 874–885.

Kumar A, Sandhu N, Dixit S, Yadav S, Swamy BPM, Shamsudin NAA. (2018) Marker-assisted selection strategy to pyramid two or more QTLs for quantitative trait-grain yield under drought. Rice 11: 35.

Liu C, Mao B, Ou S, Wang W, Liu L, Wu Y, Chu C, Wang X. (2014) OsbZIP71, a bZIP transcription factor, confers salinity and drought tolerance in rice. Plant Molecular Biology 84: 19–36.

Liu C, Ou S, Mao B, Tang J, Wang W, Wang H, Cao S, Schlappi MR, Zhao B, Xiao G, et al. (2018) Early selection of bZIP73 facilitated adaptation of japonica rice to cold climates. Nature Communication 9: 3302.

Liu H, Dong S, Li M, Gu F, Yang G, Guo T, Chen Z, Wang J. (2021) The Class III peroxidase gene OsPrx30, transcriptionally modulated by the AT-hook protein OsATH1, mediates rice bacterial blight-induced ROS accumulation. Journal of Integrative Plant Biology 63: 393–408.

Ma X, Zhang Q, Zhu Q, Liu W, Chen Y, Qiu R, Wang B, Yang Z, Li H, Lin Y, et al. (2015) Arobust CRISPR/Cas9 system for convenient, high-efficiency multiplex genome editing in monocot and dicot plants. Molecular Plant 8: 1274–1284.

Mao C, Lu S, Lv B, Zhang B, Shen J, He J, Luo L, Xi D, Chen X, Ming F. (2017) Arice NAC transcription factor promotes leaf senescence via ABA biosynthesis. Plant Physiology 174: 1747–1763.

Miao J, Li X, Li X, Tan W, You A, Wu S, Tao Y, Chen C, Wang J, Zhang D, et al. (2020) OsPP2C09, a negative regulatory factor in abscisic acid signalling, plays an essential role in balancing plant growth and drought tolerance in rice. New Phytologist 227: 1417–1433.

Min MK, Choi EH, Kim JA, Yoon IS, Han S, Lee Y, Lee S, Kim BG. (2019) Two clade Aphosphatase 2Cs expressed in guard cells physically interact with abscisic acid signaling components to induce stomatal closure in rice. Rice 12: 37.

Mohsenifard E, Ghabooli M, Mehri N, Bakhshi BB. (2017) Regulation of miR159 and miR396 mediated by Piriformospora indica confer drought tolerance in rice. Journal of Plant Molecular Breeding. 5: 10–18.

Nadarajah K, Kumar IS. (2019) Drought response in rice: the miRNA story. International Journal of Molecular Sciences 20: 3766.

Nambara E, Marion-Poll A. (2005) Abscisic acid biosynthesis and catabolism. Annual Review of Plant Biology 56: 165–185.

Nantel A, Quatrano RS. (1996) Characterization of three rice basic/leucine zipper factors, including two inhibitors of EmBP-1 DNA binding activity. Journal of Biological Chemistry 271: 31296–31305.

Nijhawan A, Jain M, Tyagi AK, Khurana JP. (2008) Genomic survey and gene expression analysis of the basic leucine zipper transcription factor family in rice. Plant Physiology 146: 333–350.

Nonhebel HM, Griffin K. (2020) Production and roles of IAA and ABA during development of superior and inferior rice grains. Functional Plant Biology 47: 716–726.

Rehman A, Azhar MT, Hinze L, Qayyum A, Li H, Peng Z, Qin G, Jia Y, Pan Z, He S, et al. (2021) Insight into abscisic acid perception and signaling to increase plant tolerance to abiotic stress. Journal of Plant Interactions 16: 222–237.

Tang N, Zhang H, Li X, Xiao J, Xiong L. (2012) Constitutive activation of transcription factor OsbZIP46 improves drought tolerance in rice. Plant Physiology 158: 1755–1768.

Wang Y, Hou Y, Qiu J, Wang H, Wang S, Tang L, Tong X, Zhang J. (2020) Abscisic acid promotes jasmonic acid biosynthesis via a ‘SAPK10-bZIP72-AOC’ pathway to synergistically inhibit seed germination in rice (Oryza sativa). New Phytologist 228: 1336–1353.

Wang Z, Chen C, Xu Y, Jiang R, Han Y, Chong K. (2004) Apractical vector for efficient knockdown of gene expression in rice (Oryza sativa L.). Plant Molecular Biology Reporter 22: 409–417.

Xia K, Ou X, Tang H, Wang R, Wu P, Jia Y, Wei X, Xu X, Kang SH, Kim SK, et al. (2015a) Rice microRNA osa-miR1848 targets the obtusifoliol 14alpha-demethylase gene OsCYP51G3 and mediates the biosynthesis of phytosterols and brassinosteroids during development and in response to stress. New Phytologist 208: 790–802.

Xia K, Ou X, Gao C, Tang H, Jia Y, Deng R, Xu X, Zhang M. (2015b) OsWS1 involved in cuticular wax biosynthesis is regulated by osa-miR1848. Plant, Cell & Environment 38: 2662–2673.

Xiang Y, Tang N, Du H, Ye HY, Xiong LZ. (2008) Characterization of OsbZIP23 as a key player of the basic leucine zipper transcription factor family for conferring abscisic acid sensitivity and salinity and drought tolerance in rice. Plant Physiology 148: 1938–1952.

Xue LJ, Zhang JJ, Xue HW. (2009) Characterization and expression profiles of miRNAs in rice seeds. Nucleic Acids Research 37: 916–930.

Yan J, Gu Y, Jia X, Kang W, Pan S, Tang X, Chen X, Tang G. (2012) Effective small RNAdestruction by the expression of a short tandem target mimic in Arabidopsis. Plant Cell 24: 415–427.

Yan J, Zhao CZ, Zhou JP, Yang Y, Wang PC, Zhu XH, Tang GL, Bressan RA, Zhu JK. (2016) The miR165/166 mediated regulatory module plays critical roles in ABA homeostasis and response in Arabidopsis thaliana. PLoS Genetics 12: e1006416.

Ye N, Zhu G, Liu Y, Li Y, Zhang J. (2011) ABA controls H2O2 accumulation through the induction of OsCATB in rice leaves under water stress. Plant & Cell Physiology 52: 689–698.

Yi R, Zhu ZX, Hu JH, Qian Q, Dai JC, Ding Y. (2013) Identification and expression analysis of microRNAs at the grain filling stage in rice (Oryza sativa L.) via deep sequencing. PLoS One 8: e57863.

Zhang F, Xiang L, Yu Q, Zhang H, Zhang T, Zeng J, Geng C, Li L, Fu X, Shen Q, et al. (2018) ARTEMISININ BIOSYNTHESIS PROMOTING KINASE 1 positively regulates artemisinin biosynthesis through phosphorylating AabZIP1. Journal of Experimental Botany 69: 1109–1123.

Zhou L, Liu Y, Liu Z, Kong D, Luo L. (2010) Genome-wide identification and analysis of drought-responsive microRNAs in Oryza sativa. Journal of Experimental Botany 61: 4157–4168.

Zhu G, Ye N, Zhang J. (2009) Glucose-induced delay of seed germination in rice is mediated by the suppression of ABA catabolism rather than an enhancement of ABA biosynthesis. Plant Cell Physiology 50: 644–651.

Zong W, Tang N, Yang J, Peng L, Ma S, Xu Y, Li G, Xiong L. (2016) Feedback Regulation of ABA signaling and biosynthesis by a bZIP transcription factor targets drought-resistance-related genes. Plant Physiology 171: 2810–2825.

